# Measurement Method Influences the Interpreted Effect of Oral Gavage on Murine Circadian Activity

**DOI:** 10.64898/2026.04.10.717555

**Authors:** Brooke A Prakash, Guohao Ni, Aarti Jagannath, Sridhar R Vasudevan

## Abstract

Historically, the primary method for measuring murine circadian activity *in vivo* has been monitoring voluntary wheel running. Recently, passive infrared (PIR) motion sensors have emerged as an alternative that is not reliant on voluntary behaviour. While research has examined the differences between the two methods for measuring circadian parameters, little focus has been placed on how these techniques may confound the assessment of therapeutic interventions. Here, we show that wheel running activity is disproportionately affected by daily oral gavage of saline compared to sham gavage treatment. In contrast, PIR-monitored activity indicates little difference between the two treatments. Both PIR and running-wheel-measured activity show a reduction in circadian amplitude and an increase in intradaily variability during both types of gavage, likely reflecting the stress of daily gavage, though the mice showed no weight loss. This finding indicates that pre- and post-intervention comparisons will misattribute gavage effects to the intervention unless appropriate sham and vehicle controls are included. More broadly, the choice of circadian measurement technique fundamentally shapes the interpretation of pharmacological interventions and must be considered in experimental design.

## Introduction

Mammalian physiology and behaviour are temporally organised by an endogenous circadian clock^1^. In murine models, the primary behavioural output used to study the state of the circadian pacemaker, the suprachiasmatic nucleus, is locomotor activity. Whilst a number of technologies have been adapted to monitor locomotor activity in murine models, the most prevalent are voluntary running wheels and passive infrared (PIR) sensors. Critically, the choice between active and passive monitoring shapes the circadian phenotype observed in mice^2^. Running wheels have been widely used for circadian monitoring due to the robust and high-amplitude rhythms they produce. However, recent studies increasingly recognise that running wheels can act as non-photic zeitgebers, providing feedback to the central clock^3^. Running wheel exercise shortens the endogenous period and shifts core circadian clock gene phase in peripheral tissues, such as the liver and skeletal muscles, compared to non-wheel conditions^4^. In contrast, while PIR data is traditionally considered noisier, it provides a passive way to monitor home cage activity by capturing grooming and feeding behaviour in mice^5^ Thus, it allows behavioural recording without the confounding influence of exercise-induced feedback.

When studying circadian pharmacology and investigating the influence of certain drugs on circadian rhythms, an often-overlooked variable is the impact of the delivery method on behavioural output. For example, oral gavage, a common method of precision drug delivery, can induce a measurable stress response, significantly increasing plasma corticosterone levels following repeated interventions and altering gastrointestinal function^6^. The activation of the hypothalamic-pituitary-adrenal axis via repeated oral gavage can be problematic for circadian research, as glucocorticoid signalling can directly interfere with peripheral clock gene expression, leading to masking effects that mimic a clock shift but are, in reality, acute responses to stress or pain^7^. Therefore when assessing the effects of drugs on circadian rhythms, the method of drug administration must be carefully considered. Some experimental designs compare a baseline period of activity with a post-treatment period of activity^8^. This approach risks conflating gavage effects with the drug’s true effect unless proper controls are included^9^. Sham-control groups, in which mice are familiarised with the gavage procedure prior to drug administration, are therefore recommended^10^.

Here we compare the sensitivity of running wheel and PIR systems in monitoring circadian disruptions caused by sham and saline oral gavage treatments in murine models. Such methodological comparisons are essential for ensuring that observed effects reflect pharmacological activity rather than artefacts of experimental design.

## Methods

### Animals

Male C57BL/6J mice were purchased from Charles River and individually housed under a 12:12 Light-Dark (LD) cycle with access to food and water *ad libitum*. Data collection began at 19 weeks old. This study was restricted to male mice as the oestrous cycle produces scalloped circadian rhythms, making assessment of rhythmicity difficult over the short term in females. All procedures were performed in accordance with the UK Home Office Animals (Scientific Procedures) Act 1986 and the University of Oxford’s Policy on the Use of Animals in Scientific Research.

### Wheel-running Circadian Monitoring

Mice were individually housed in cages with wheels. Voluntary wheel-running activity was recorded via wheel-rotation counts collected using equipment from Actimetrics (Wilmette, IL).

### Passive Infrared Circadian Monitoring

Mice were individually housed under passive infrared (PIR) motion sensors following previously published methods^5.^ Data collection and storage was carried out using Processing 4.3 (Cambridge, MA).

### Oral Gavage

Animals were gavaged daily with saline (10mL/kg) or sham (no liquid) 30 minutes before lights off (ZT 12) for four days at a time.

### Circadian Analyses

All wheel-running and PIR data was visualised and analysed in Clocklab (Actimetrics, Wilmette, IL). Amplitude, intradaily variability (IV), and interdaily stability (IS) were calculated over a 4-day period.

### Statistical Analysis

GraphPad Prism 10 (Boston, MA) was used to perform statistical analyses as described in figure legends. All data are represented as individual replicates and mean ± SEM. In all figures,*= P ≤ 0.05, ** = P ≤ 0.01, *** = P ≤ 0.001.

## Results

To separate the effect of gavage itself from liquid insertion, we serially treated five mice with saline followed by sham gavage (Fig. 1). These five mice were acclimatised in singly housed cages with running wheels. At 20 weeks, they were administered saline (10ml/kg) via oral gavage at zeitgeber time (ZT) 11.5 for four consecutive days while voluntary wheel-running activity was recorded. After three days of recovery the same mice were treated with daily sham gavage at ZT 11.5 for four days to allow internal comparison within the same mouse.

**Fig. 1.**
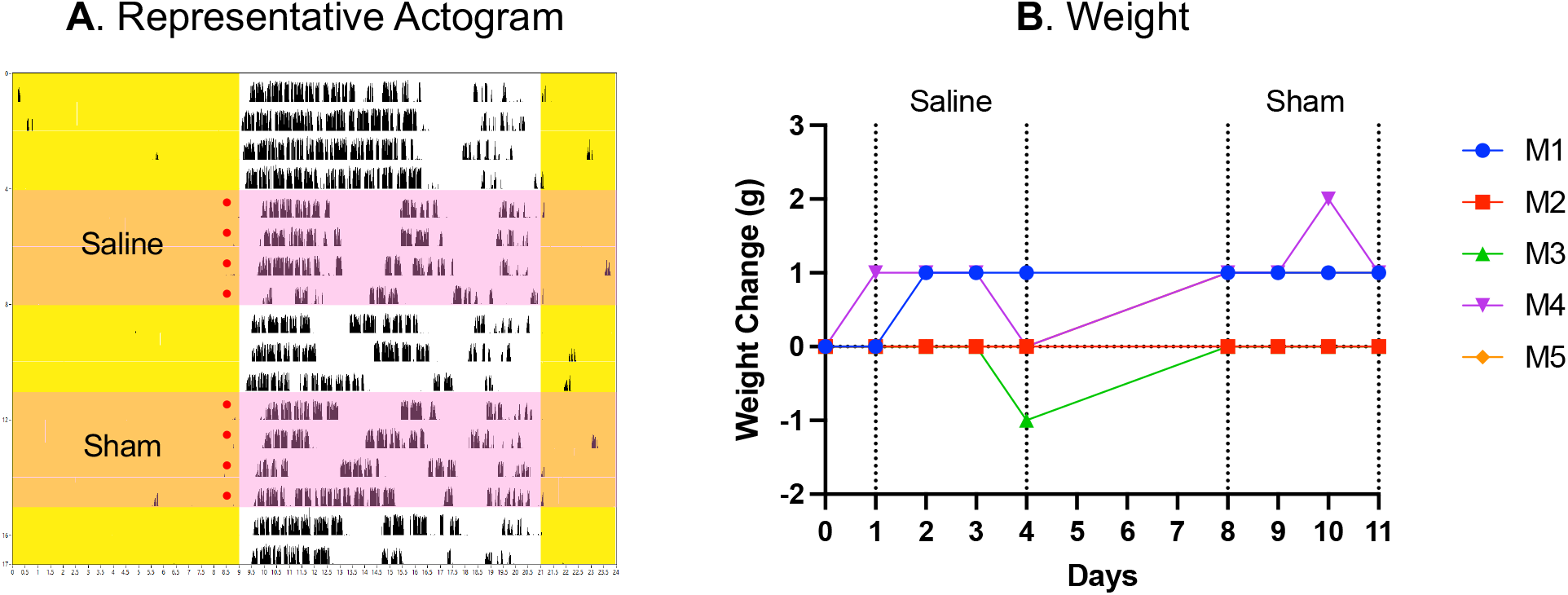
Both saline and sham gavage alter mouse wheel-running activity. **(A)** Representative actogram of an animal receiving saline via oral gavage for four days, followed by three days of recovery and then sham treatment for four days. Yellow denotes lights on, light pink denotes period of gavage treatment, red dot denotes time of daily gavage. **(B)** Weight change in grams over the course of gavaging mice (M1-M5) on running wheels.

Both saline and sham gavage altered wheel-running activity without causing prolonged weight loss (Fig. 1). We compared baseline activity of four days before gavage treatment with four days of saline gavage treatment and four days of sham gavage using repeated measures one-way ANOVA. These analyses revealed that sham gavage treatment had less of an impact on running-wheel activity than saline gavage treatment, though as expected the process of gavage itself significantly impacted wheel-running activity (Fig. 2). Gavage treatment significantly reduced the strength (amplitude) of voluntary wheel-running activity (Fig. 2A), increased the variability of daily activity (Fig. 2B) and decreased the stability of circadian activity (Fig. 2C). The effect was less pronounced during sham gavage specifically for amplitude and intradaily variability (IV). Both saline and sham gavage significantly reduced mean hourly wheel-running activity compared to baseline. However, the animals showed a relative recovery of activity at four of the six reduced timepoints when sham treated (Fig. 2D).

**Fig. 2.**
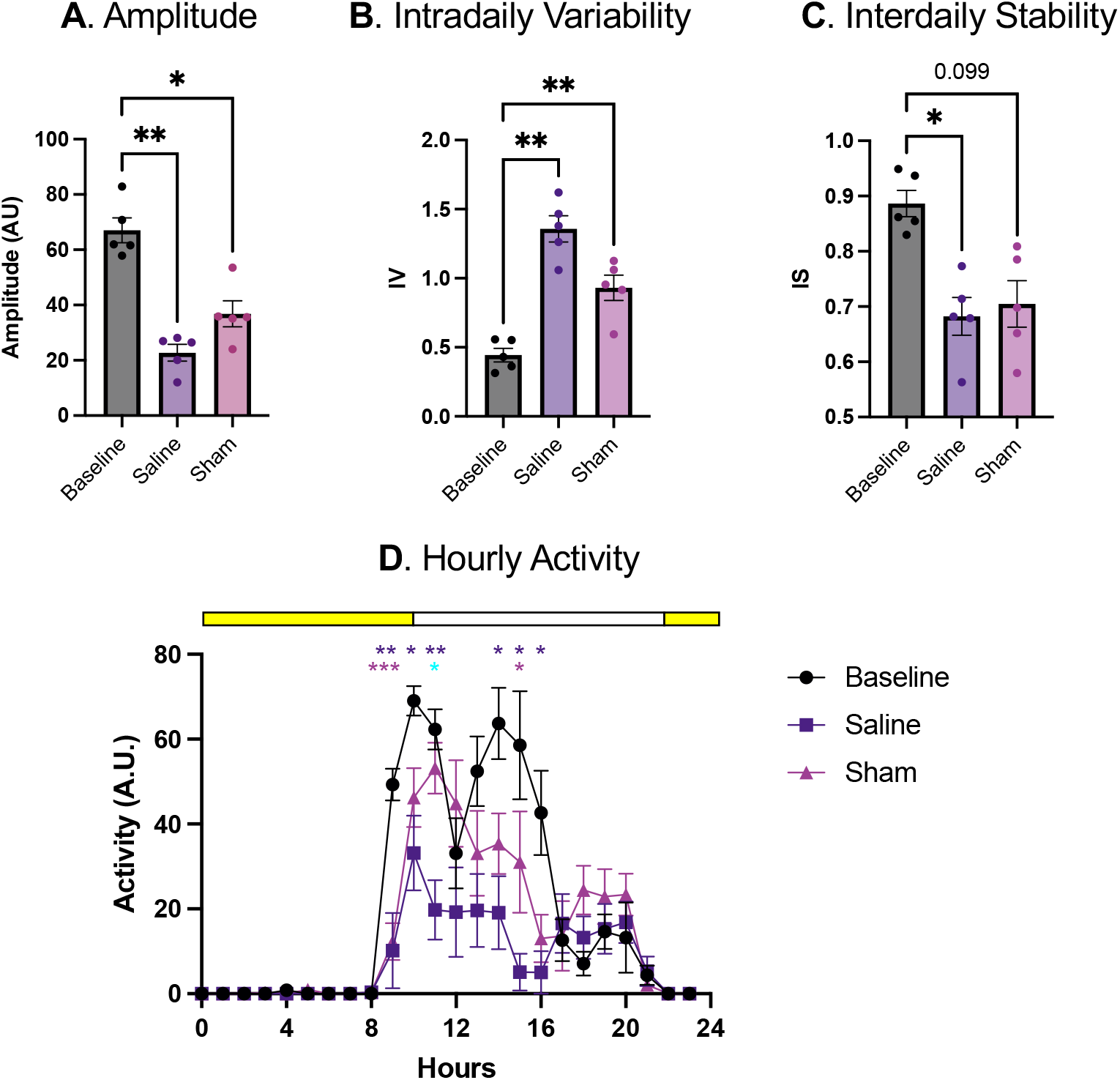
Oral gavage alters circadian parameters measured via wheel-running activity. Circadian **(A)** amplitude, **(B)** intradaily variability, and **(C)** interdaily stability were calculated over four-day periods before treatment (baseline), during daily saline gavage, and during daily sham gavage. Data analysed with repeated measures one-way ANOVA with Tukey’s multiple comparisons test, N=5. P=0.0008 and F(1.788, 7.153) = 23.34 for **A**, P=0.0033 and F(1.283, 5.130) = 24.60 for **B**, P=0.0174 and F(1.704, 6.816) = 8.165 for **C.(D)** Mean activity per hour was calculated for each mouse on the last day of each treatment indicated. Light bar indicates lights on/off. Colour of star indicates significant difference from baseline, except blue which indicates a significant difference between saline and sham. Data analysed with repeated measures two-way ANOVA with Tukey’s multiple comparisons test, N=5. P<0.0001 and F(2.589, 10.36) = 22.99 for time, P=0.0047, and F(1.191, 4.763) = 23.64 for treatment, and P=0.0183 and F(3.035, 12.14) = 4.916 for time x treatment.

Given that these measures were calculated from wheel-running, a voluntary activity that mice do not perform continuously during waking periods, we used PIR monitoring to evaluate whether gavage effects on circadian parameters differed across measurement techniques. We therefore moved these mice into single-housing cages monitored with PIR motion sensors and added an additional seven age- and experience-matched mice for group comparisons. Circadian analyses compared four days of baseline activity to four days of activity during daily saline (N=6) or sham (N=6) gavages at ZT 11.5. This time data was analysed with repeated measures two-way ANOVA, since there were two separate groups receiving sham and saline treatment.

Once again, the process of gavage itself altered measured activity in both saline and sham conditions without inducing weight loss (Fig. 3). There was no significant difference between the circadian amplitude of sham and saline gavage treatments, but as expected both treatments significantly reduced amplitude (Fig. 4A). Meanwhile changes in IV demonstrated a non-significant increase in instability in both conditions (Fig. 4B) while IS was only significantly reduced for the saline-treated group (Fig. 4C). The decrease in mean hourly activity was only significant for one hour post-gavage, indicating that the reduction of activity was more transient when measured with PIR (Fig. 4D).

**Fig. 3.**
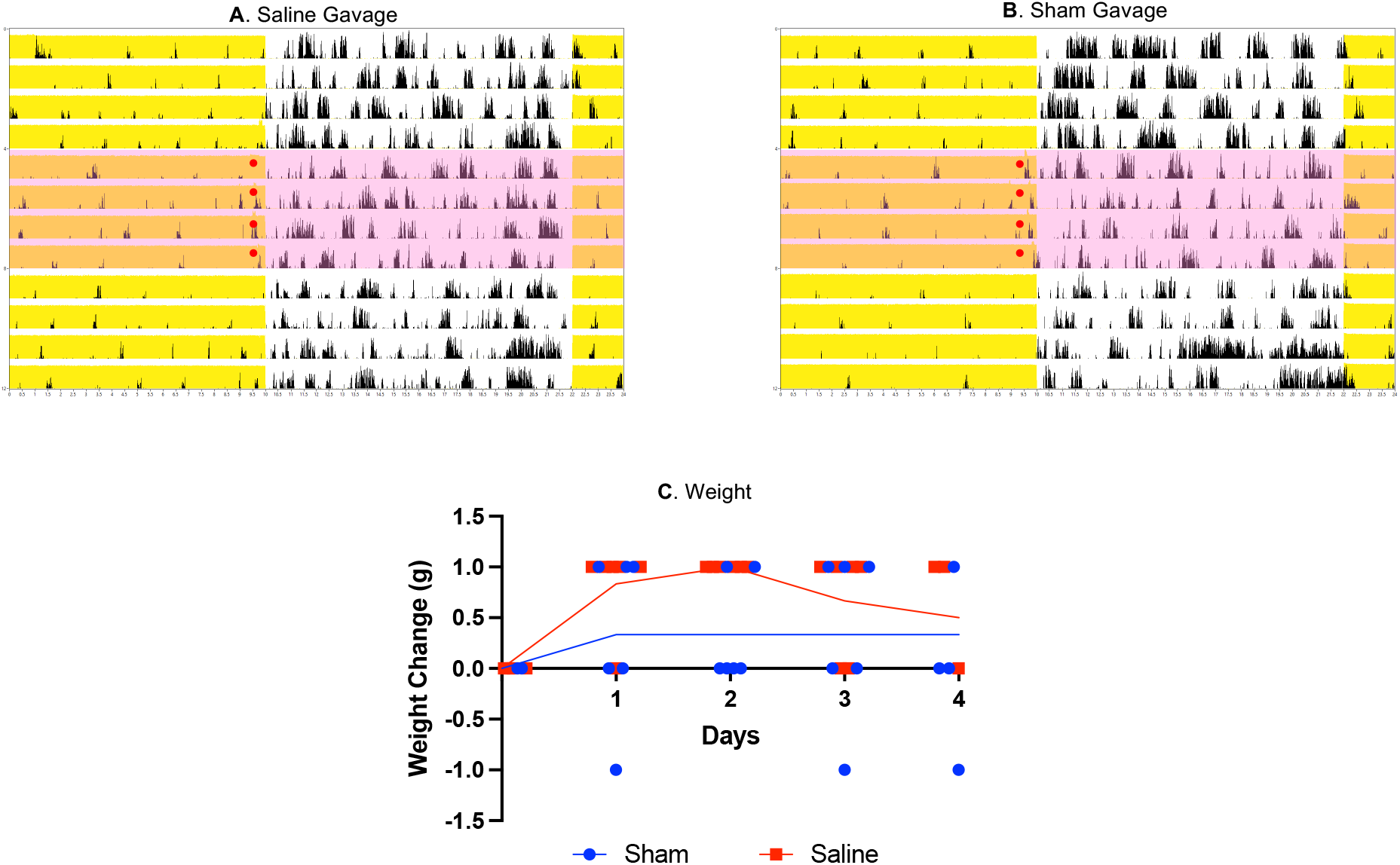
Both saline and sham gavage alter mouse activity recorded with PIR. Representative actograms of **(A)** saline and **(B)** sham-treated animals recorded with PIR. Yellow denotes lights on, light pink denotes period of gavage treatment, red dot denotes time of daily gavage. **(C)** Weight change in grams over the course of gavaging mice during PIR monitoring. Mean with individual weights plotted, N=6.

**Fig. 4.**
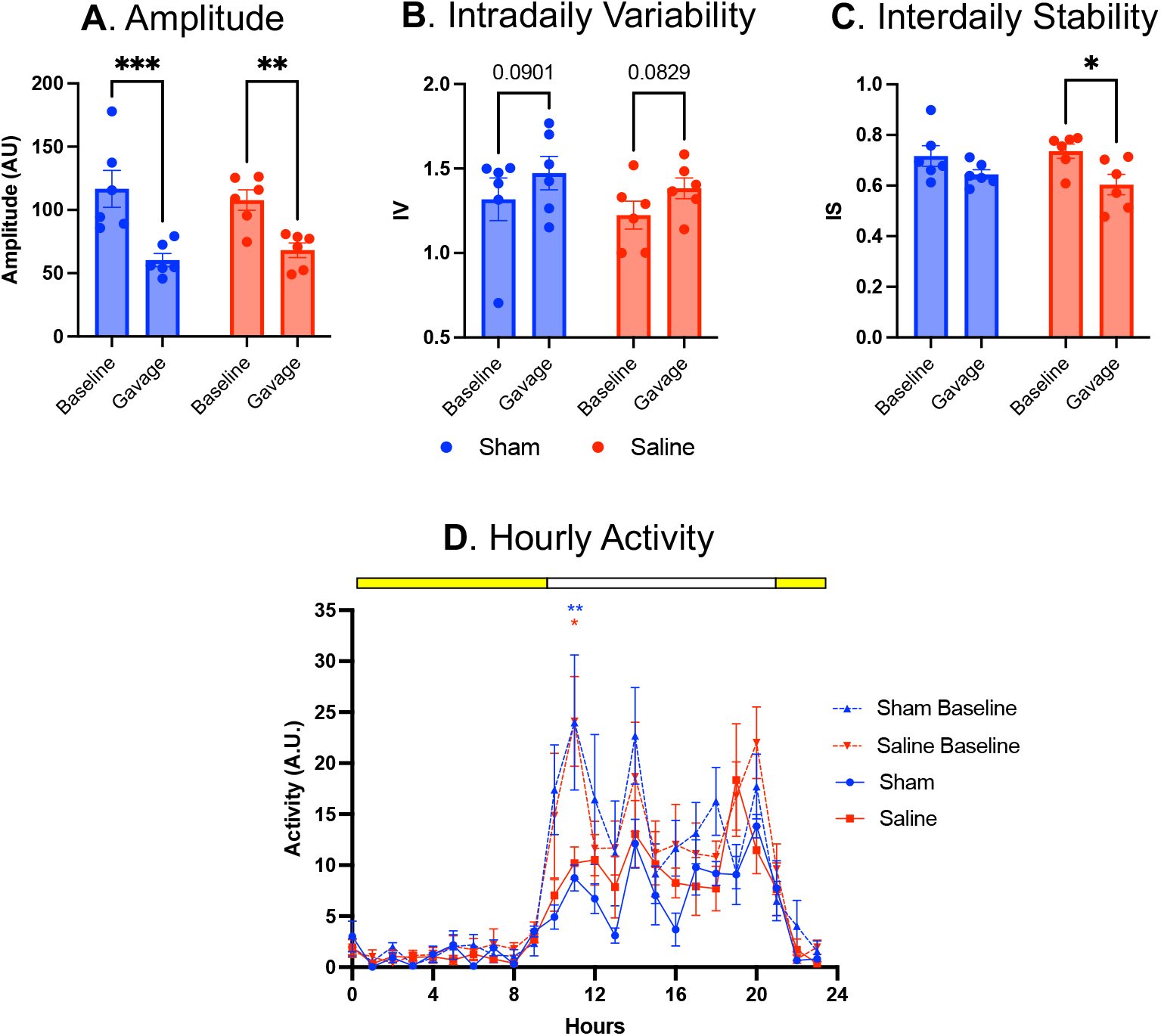
Oral gavage alters circadian parameters measured via PIR activity. Circadian **(A)** amplitude, **(B)** intradaily variability, and **(C)** interdaily stability were calculated over four-day periods before treatment (baseline) and during daily saline or sham gavages. Data analysed with repeated measures two-way ANOVA followed by uncorrected Fisher’s LSD with single pooled variance, N=6. P<0.0001 and F(1.191, 4.763) = 23.64 for gavage in **A**, P=0.0227 and F(1, 10) = 7.230 for gavage in **B**, P=0.0075 and F(1, 10) = 11.15 for gavage in **C**. P=nonsignificant for gavage type in **A**,**B**,**C. (D)** Mean activity per hour was calculated for each mouse on the last day of each treatment indicated. Light bar indicates lights on/off. Colour of star indicates significant difference from baseline. Data analysed with repeated measures three-way ANOVA with Tukey’s multiple comparisons test as sham and saline-treated animals were separate groups, N=6. P<0.0001 and F(23, 230) = 21.44 for time, P=0.0015 and F(1, 10) = 18.64 for gavage, P<0.0001 and F(23, 230) = 3.281 for time x gavage and P=nonsignificant for gavage type.

To directly compare the measures collected with running-wheel activity to the measures collected via PIR, we normalised the parameters collected in each to its baseline (Fig. 5). Both running-wheel and PIR activity showed a general reduction in amplitude and increase in IV during daily gavages. Running-wheel activity, however, showed a pronounced reduction in circadian amplitude during saline gavage (Fig. 5A). IV was significantly lower when estimated by PIR than by wheel running. This result suggests that running-wheel variability is more impacted by the stress of gavage itself. This effect was especially stark in the saline-treated wheel-running group (Fig. 5B). The only metric that was unaffected by measurement technique was IS, which was estimated similarly by PIR and wheel-running across groups (Fig. 5C).

**Fig. 5.**
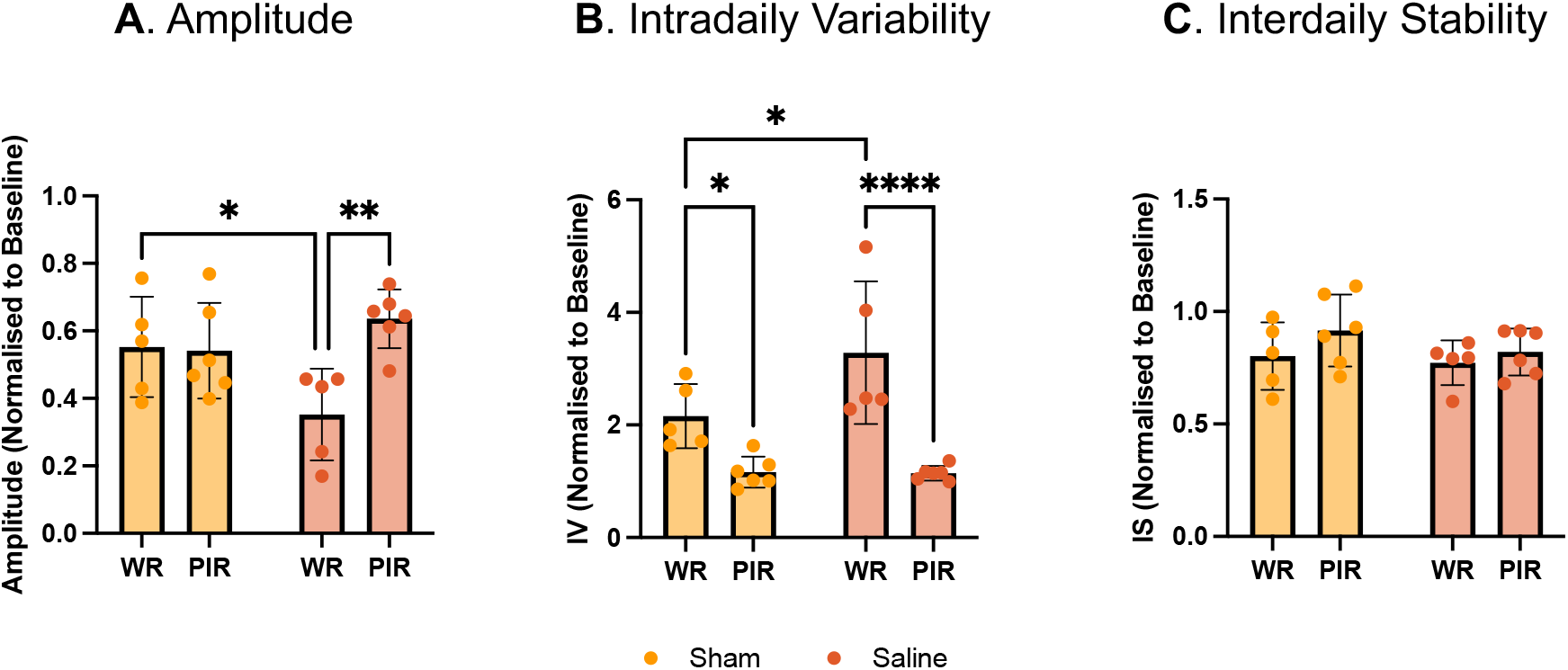
Measurement technique influences circadian parameter estimation during oral gavage. Circadian **(A)** amplitude, **(B)** intradaily variability, and **(C)** interdaily stability were calculated over four-day periods before treatment (baseline) and during daily saline or sham gavages. Data analysed with two-way ANOVA followed by uncorrected Fisher’s LSD with single pooled variance, N=5-6. P=0.0240 and F(1, 18) = 6.072 for measurement method, P=0.0158 and F(1, 18) = 7.096 for measurement method x gavage type in **A**, P<0.0001 and F(1, 18) = 29.46 for measurement method in **B**, no significant effects in **C**.

Overall, PIR measurements showed minimal differences between sham and saline-treated animals, while running-wheel measurements revealed a pronounced difference.

## Discussion

Little attention has been paid to how the choice of measurement method may alter the interpretation of therapeutic interventions. Our data clearly demonstrate that the technique used to measure circadian activity *in vivo* can fundamentally shape the interpretation of pharmacological interventions.

Saline has not been shown to behaviourally alter circadian rhythms *in vivo* besides hypertonic saline advancing the onset of activity if administered at ZT 19^11^. Our *in vivo* running-wheel activity data, however, indicates that saline significantly reduces circadian amplitude and increases intradaily variability beyond the effects of sham gavage. PIR, a measure that is not reliant on voluntary activity, showed no such distinction; sham and saline gavages were relatively indistinguishable. It is possible that running activity may be more sensitive to the intragastric volume of liquid delivered, whereas PIR captures broader home cage movement that is less affected.

Nevertheless, both running-wheel and PIR measurements revealed a reduction in circadian amplitude and an increase in intradaily variability upon the introduction of daily gavage – whether sham or not. One might suggest these effects are transient, given the four-day treatment window; however, as others have shown a prolonged reduction in PIR activity after 25 days of daily s.c. saline injections ^12^, this seems unlikely. It is more likely that daily gavage introduced stress, altering circadian activity^13^. However, any anxiety experienced was not strong enough to cause weight loss - a commonly used indicator of stress.

Our findings have widespread implications for the design of studies on pharmacological interventions in murine circadian rhythms *in vivo*. For example, pre- and post-treatment comparisons will misattribute gavage effects to the intervention unless a sham or vehicle control is included. PIR captures a broader range of behaviours, making rhythms appear less robust and noisier. Wheel-running offers cleaner readouts of activity and can be preferable depending on the experimental context. Based on our results, we therefore suggest that a vehicle (rather than a sham) control is compared to a treated mouse if using wheel-running activity to estimate the effect of a therapeutic intervention administered by oral gavage.

One limitation of the current study is that it only assesses one method of drug administration. Future research should compare the impact of daily oral, intraperitoneal, subcutaneous, or food administration across different measures of circadian activity to characterise the effects of each. Such work will help distinguish true pharmacological effects on circadian activity from artefacts of the administration procedure. As circadian pharmacology advances towards clinical translation, methodological rigour in activity monitoring will be essential for accurate interpretation of therapeutic effects.

## Acknowledgements

The authors thank the University of Oxford Biomedical Services for animal husbandry support. This work was supported by the Michael J. Fox Foundation for Parkinson’s Research (grant MJFF-025202). The authors declare no conflicts of interest.

## References

1. Czeisler CA, Duffy JF, Shanahan TL, Brown EN, Mitchell JF, Rimmer DW et al. Stability, precision, and near-24-hour period of the human circadian pacemaker. Science 1999; 284(5423): 2177–2181.

2. Brown LA, Banks GT, Horner N, Wilcox SL, Nolan PM, Peirson SN. Simultaneous Assessment of Circadian Rhythms and Sleep in Mice Using Passive Infrared Sensors: A User’s Guide. Curr Protoc Mouse Biol 2020; 10(3): e81.

3. Oneda S, Cao S, Haraguchi A, Sasaki H, Shibata S. Wheel-Running Facilitates Phase Advances in Locomotor and Peripheral Circadian Rhythm in Social Jet Lag Model Mice. Front Physiol 2022; 13: 821199.

4. Yasumoto Y, Nakao R, Oishi K. Free access to a running-wheel advances the phase of behavioral and physiological circadian rhythms and peripheral molecular clocks in mice. PLoS One 2015; 10(1): e0116476.

5. Brown LA, Hasan S, Foster RG, Peirson SN. COMPASS: Continuous Open Mouse Phenotyping of Activity and Sleep Status. Wellcome Open Res 2016; 1: 2.

6. Hanning N, Verboven R, De Man JG, Ceuleers H, De Schepper HU, Smet A et al. Single-day and multi-day exposure to orogastric gavages does not affect intestinal barrier function in mice. Am J Physiol Gastrointest Liver Physiol 2023; 324(4): G281–G294.

7. Koch CE, Leinweber B, Drengberg BC, Blaum C, Oster H. Interaction between circadian rhythms and stress. Neurobiol Stress 2017; 6: 57–67.

8. Brenna A, Ripperger JA, Albrecht U. Locomotor Activity Monitoring in Mice to Study the Phase Shift of Circadian Rhythms Using ClockLab (Actimetrics). Bio Protoc 2025; 15(4): e5187.

9. Walker MK, Boberg JR, Walsh MT, Wolf V, Trujillo A, Duke MS et al. A less stressful alternative to oral gavage for pharmacological and toxicological studies in mice. Toxicol Appl Pharmacol 2012; 260(1): 65–69.

10. Karrberg L, Andersson L, Kastenmayer RJ, Ploj K. Refinement of habituation procedures in diet-induced obese mice. Lab Anim 2016; 50(5): 397–399.

11. Gizowski C, Bourque CW. Sodium regulates clock time and output via an excitatory GABAergic pathway. Nature 2020; 583(7816): 421–424.

12. Collins HM, Pinacho R, Tam SKE, Sharp T, Bannerman DM, Peirson SN. Continuous home cage monitoring of activity and sleep in mice during repeated paroxetine treatment and discontinuation. Psychopharmacology (Berl) 2023; 240(11): 2403–2418.

13. Turner PV, Brabb T, Pekow C, Vasbinder MA. Administration of substances to laboratory animals: routes of administration and factors to consider. J Am Assoc Lab Anim Sci 2011; 50(5): 600–613.

